# Automated detection of Bornean white-bearded gibbon (*Hylobates albibarbis*) vocalisations using an open-source framework for deep learning

**DOI:** 10.1101/2024.04.15.589517

**Authors:** A. F. Owens, Kimberley J. Hockings, Muhammed Ali Imron, Shyam Madhusudhana, Mariaty, Tatang Mitra Setia, Manmohan Sharma, Siti Maimunah, F. J. F. Van Veen, Wendy M. Erb

## Abstract

Passive acoustic monitoring is a promising tool for monitoring at-risk populations of vocal species, yet extracting relevant information from large acoustic datasets can be time-consuming, creating a bottleneck at the point of analysis. To address this, we adapted an open-source framework for deep learning in bioacoustics to automatically detect Bornean white-bearded gibbon (*Hylobates albibarbis*) “great call” vocalisations in a long-term acoustic dataset from a rainforest location in Borneo. We describe the steps involved in developing this solution, including collecting audio recordings, developing training and testing datasets, training neural network models, and evaluating model performance. Our best model performed at a satisfactory level (F score = 0.87), identifying 98% of the highest-quality calls from 90 hours of manually-annotated audio recordings and greatly reduced analysis times when compared to a human observer. We found no significant difference in the temporal distribution of great call detections between the manual annotations and the model’s output. Future work should seek to apply our model to long-term acoustic datasets to understand spatiotemporal variations in H. albibarbis’ calling activity. Overall, we present a roadmap for applying deep learning to identify the vocalisations of species of interest which can be adapted for monitoring other endangered vocalising species.

## I. INTRODUCTION

Ever-increasing anthropogenic pressures on the environment, such as habitat loss, have led to widespread population declines in many animal species (Bender et al. 1998). However, for many species, data on population trends are often sparse (Jetz et al. 2019), leading to an increased demand for wildlife population monitoring programs to inform conservation responses (Verma et al. 2016). To help achieve this, conservation scientists and ecologists have turned to developing technologies to automate data collection, enabling the rapid accumulation of large volumes of data (Piel and Wich 2021). While this has allowed for unprecedented insight, it can also make practical aspects of ecological inference challenging (Borowiec et al. 2022).

Manual extraction of relevant information from large datasets can be time consuming, resulting in a bottleneck at the point of analysis (Norouzzadeh et al. 2018). This bottleneck is evident in data generated as part of passive acoustic monitoring (PAM) programs, which involve the use of autonomous acoustic sensors to collect sound recordings in the field (Acevedo and Villanueva-Rivera 2010). Advancements in recording device design and cost, as well as improved data storage options have made the task of capturing many hours of acoustic data relatively straightforward (Morgan and Braasch 2021; Piel and Wich 2021). Data must then be browsed to identify relevant signals of interest, such as species-specific vocalisations, often by manually listening to each recording in full or visually inspecting the data in spectrogram form (a time-frequency pictorial representation of an audio signal), or both (van Kuijk et al. 2023; Clink et al. 2023). PAM can provide a step-change in standardized population monitoring of vocal species at high temporal resolution and simultaneous large spatial scales, which would be impossible to achieve with ‘traditional’ methods relying on manual data collection (Sugai et al. 2019). However, such PAM programs often capture datasets so large that they cannot be studied manually in full in a reasonable timeframe, so automating this limiting data-processing step is critical (Morgan and Braasch 2021; Clink et al. 2023).

Machine learning has proven to be an effective solution for fast and accurate analysis of acoustic data, including the automated detection of signals of interest (Stowell 2022; Miller et al. 2023). There are many options available for this task, including artificial neural networks (ANNs) (Mielke and Zuberbühler 2013), Gaussian mixture models (GMMs) (Heinicke et al. 2015) and support vector machines (SVMs) (Noda et al. 2016), among others (reviewed in Knight et al. 2017). These each have associated advantages, and due to the diversity of potential signal types and acoustic environments, no single method is optimal in all situations (Clink et al. 2023). However, it is worth noting that ANNs demonstrate comparatively strong adaptability and proficiency in understanding complex patterns in data (Haykin 2009; Bengio et al. 2016). ANNs can be arranged in various architectures, each suited for different tasks. Early work applying ANNs to animal sound made use of the multi-layer perception (MLP) architecture, with manually selected summary features such as syllable duration, peak frequency, etc., used to inform the network’s predictions (Stowell 2022). While MLPs have been effective in classifying a wide variety of terrestrial and marine animal calls, the structure of non-speech acoustic events can be highly variable (Kong et al. 2017), and reducing the data to a series of manually assigned summary features can restrict the wealth of information available to train a network, potentially limiting its effectiveness (Stowell 2022).

Deep neural networks, such as Convolutional Neural Network (CNN) architectures, rely on feature sets that are not manually selected, but instead learned during the training process (Morgan and Braasch 2021). CNNs are particularly effective for processing visual representations of audio, such as spectrograms, leveraging their ability to learn patterns that occur both spatially and temporally in data. This allows CNNs to learn local features regardless of their spatial position within an image (Knight et al. 2017). CNNs are therefore ideal candidates for the automated detection of signals within bioacoustic data, where instances of relevant features within a spectrogram are not predefined or readily identifiable (Stowell 2022). They have been used to analyse vocalisations from a variety of taxa, including insects (Hibino et al. 2021), fish (Guyot et al. 2021), anurans (Colonna et al. 2016), birds (Narasimhan et al. 2017), marine mammals (Miller et al. 2023), bats (Mac Aodha et al. 2018), and other terrestrial mammals (Bjorck et al. 2019), including primates (Wood et al. 2023).

The potential of CNNs is far from fully realized however (Rammer and Seidl 2019), and there are relatively few examples of CNNs being used to answer well-defined research questions in ecology, as is so with other deep learning approaches (Dufourq et al. 2021). Additionally, there are few guidelines on how to approach key steps such as model tuning and performance assessments (Knight et al. 2017; Patterson and Gibson 2017; Stowell 2022). Further case studies reporting successful applications will advance the development of best practices for overcoming these challenges (Dufourq et al. 2021).

Gibbons (family Hylobatidae) are ideal candidates for the automated detection of species-specific vocalisations. They engage in loud, highly stereotyped song bouts which are largely confined to a few-hour window before and after sunrise (Cheyne et al. 2008). During a particular calling bout, they usually emit multiple calls, which facilitates the generation of abundant training data (Clink et al. 2023). Since the great call is performed largely by mated females, it is often used as an indicator of a gibbon family group, allowing for group density and spatial distribution estimates to be derived from great call densities, assuming that estimates of female calling rates are available (Cheyne et al. 2016). Furthermore, gibbons reside exclusively in tropical forests, which are often visually challenging and inaccessible so studying their populations using visual methods, such as line transect and camera trap surveys, is typically very difficult (Vu and Tran 2019). For these reasons, gibbons are model organisms for developing and testing guidelines for automated detection. So far, this has been done for a handful of species, including the Hainan gibbon (*Nomascus hainanus*) (Dufourq et al. 2021), Western black-crested gibbon (*Nomascus concolor*) (Zhou et al. 2023), Northern grey gibbon (*Hylobates funereus*) (Clink et al. 2023), and southern yellow-cheeked crested gibbon (*Nomascus gabriellae*) (Clink et al. 2024).

Here, we apply a variation of a pre-defined CNN architecture, DenseNet (Huang et al. 2016), to identify female great call vocalisations of the endangered Bornean white-bearded gibbon (*Hylobates albibarbis*) from a long-term acoustic dataset. We train a detector with high precision that minimizes false-positive rates (i.e., the rate of detections incorrectly labelled as gibbon great calls) for application to large acoustic datasets recorded by PAM arrays. This can then be used to facilitate accurate population monitoring of wild gibbons on an ever-greater spatiotemporal scale and applied as a case study for developing automated detectors for other endangered vocalising species.

## II. METHODS

### A. Data collection

The long-term acoustic dataset used in this study derives from the Mungku Baru Education and Research Forest (MBERF), a ∼50 km^2^ area of tropical rainforest in Central Kalimantan Province, Indonesia. The MBERF lies in the centre of the wider Rungan Landscape, which covers approximately 1,500 km^2^ between the Kahayan and Rungan rivers north of the provincial capital of Palangka Raya. This represents the largest area of continuous unprotected lowland rainforest remaining on the island of Borneo (Purnama and Afitah 2021). There is an estimated population of 4,000 white bearded gibbons in the Rungan Landscape and an estimated density of 2.79 groups per km^2^ in the MBERF, making the region significantly important for the conservation of the species (Buckley et al. 2018).

Eight autonomous recording units (ARUs) (Song Meter SM4, Wildlife Acoustics, Maynard, Massachusetts) were deployed here by WME in July 2018 (see Figure 1). These were placed on trees, 5 meters above the ground, in a dispersed grid with approximately 1,200 meters between each device. The MBERF contains a mosaic of different forest types, and the array was designed to capture this heterogeneity, with three ARUs situated in ‘kerangas’ (heath) forest, three in ‘low pole’ peat swamp forest, and two in ‘mixed swamp’ forest, with the latter representing a transition habitat between the former two (Buckley et al. 2018).

**FIGURE 1.**
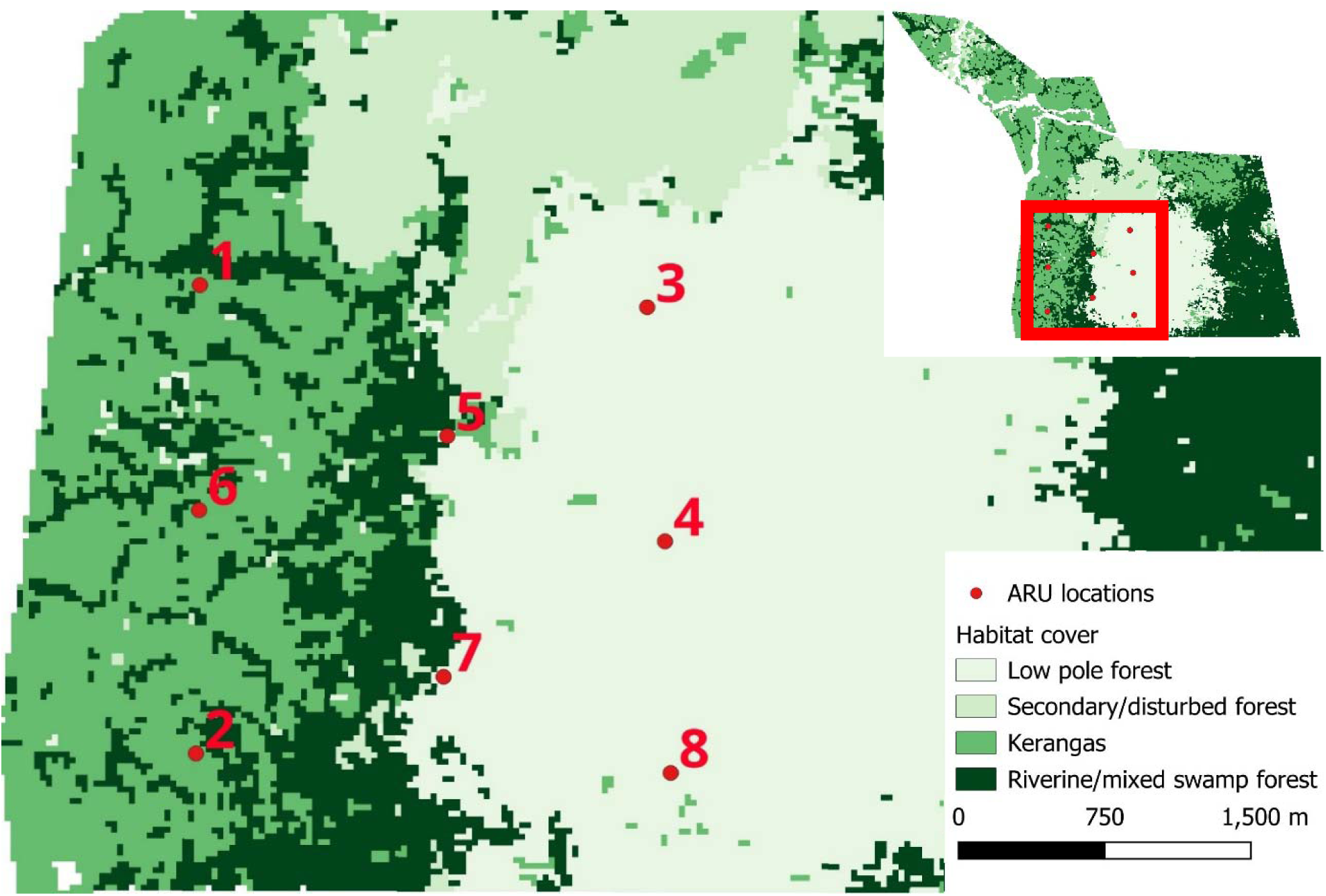
Map of the Mungku Baru Education and Research Forest (Buckley et al. 2018), showing the distribution of different habitat types over the survey area and the location of the ARUs.

The ARUs were set to record from 4 am to 6 pm (local time) daily to capture the full pre-dawn and diurnal period of ape calling. These were set to default settings (sensitivity of -35 ± 4 dB (0 dB = 1V/pa@1kHz), dynamic range of 14 to 100 dB SPL at 0 dB gain, microphone gain of 16 dB, and inbuilt preamplifier gain of 26 dB) and recorded on two channels with a sampling rate of 24 kHz. Audio was captured in 16-bit Waveform Audio File Format (WAV) and saved as one-hour files. Memory cards and batteries were changed every two weeks.

### B. Manual annotation

Manually-annotated training and testing datasets were required to develop the automated detector. To create these samples, recordings between 4-10 am were selected from a single day every four weeks from a randomly selected device for each habitat. This covers the temporal period in which most *H. albibarbis* great calls occur (Cheyne et al. 2008) and ensured that a variety of potential sound environments (i.e., capturing spatial and temporal variation) were included as training inputs to the model, improving its ability to generalize over a wider range of applications. The resultant subset contained 300 hours of recordings, covering 50 days spanning from October 2018 to December 2019.

The selected sound files were loaded into the sound analysis software Raven Pro 1.6 (K Lisa Yang Center for Conservation Bioacoustics 2024) and visualized as spectrograms using a 3462-sample Hann window with a 90% overlap and a 4096-sample Discrete Fourier Transformation. With assistance from a team of undergraduate interns (see Acknowledgements), each recording was listened to in full and visually scanned to identify instances of great calls. These were defined as vocalisations containing introductory, climax, and descending phrases (see Figure 2). A selection was created for each instance by drawing a box around the call in the spectrogram, providing information about its time-frequency boundaries.

**FIGURE 2.**
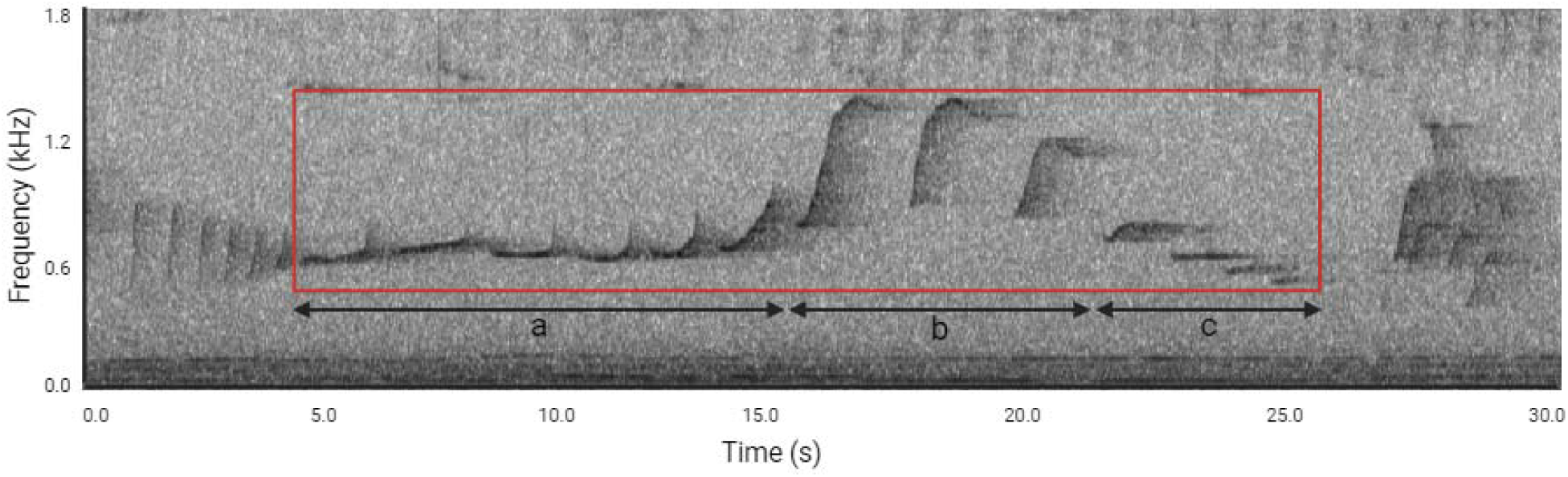
A spectrogram image of a female H. albibarbis great call, created using Raven Pro 1.6. The box represents the time and frequency boundaries of a manually annotated selection. The time boundaries span from the start of the first introductory (a) note to the end of the last descending (c) note. The frequency boundaries span from the lowest frequency descending (c) note to the highest frequency climax (b) note.

Each selection was then annotated based on its completeness and quality. A selection was marked as “clear” when the entirety of the call could be heard and was visually clear in the spectrogram, “faint” when the whole call could be heard but was not fully shown in the spectrogram (and vice versa), and very faint when the call was only partially seen and heard (i.e., part of the great call was not captured in the recording). Each selection was reviewed and edited where needed by AFO to prevent inter-observer bias. In total, 1,611 great calls were annotated.

The manually-annotated data were then randomly split, with 70% allocated for training (210 hours, 1,089 calls) and 30% allocated for testing (90 hours, 522 calls). Due to the ambiguous nature of “very faint” calls, they were removed from the training process and used solely for testing thereafter. This prevented misleading information, i.e., non-target events, being fed into the positive class weightings. Models were trained using “clear” and “faint” instances (729 calls), as well as only “clear” instances (522 calls).

### C. Automated detection

The development and testing of the automated detector utilized Koogu (version 0.7.2), an open-source framework for deep learning from bioacoustic data (Madhusudhana 2023). Koogu offers a variety of functions for deep learning, including 1) preparing audio for use as inputs to machine learning models, 2) training models, 3) assessing model performance, and 4) using trained models for the automated analysis of large datasets. For a full workflow describing how the following steps were implemented within Koogu, see Supplementary Material A.

#### 1. Data preparation

All audio files were first down-sampled to 4,500 Hz to reduce the overall file size and improve the efficiency of downstream computations (cf. Miller et al. 2023). The resulting Nyquist frequency (2,250 Hz) is above the highest frequency within the great call selections (2,077 Hz), and so no relevant information was lost in this process. The recordings were then split into consecutive 28s segments (longer than the longest manually-annotated great call at 27.7s) with a hop size of 1s, leaving an overlap of 27s between clips. The waveform of each segment was normalized by scaling the amplitudes to occur in the range -1.0 to 1.0.

The resulting start and end times of each segment were then compared to those from manual annotations. Segments that fully contained the temporal extents of an annotated great call were considered as positive inputs while segments with partial overlap were excluded from training as these could resemble non-great call events and so lead to uncertainties during the training process. Segments without temporal overlap were considered as negative inputs (i.e., background noise).

Spectrograms for both the positive and negative classes were then computed with an analysis window of 0.192s and a 75% overlap. The bandwidth was also restricted to between 200 and 2200 Hz to exclude noise outside of the target frequency range. This resulted in input spectrograms with a shape of 384 x 580 (height x width) pixels. To address the imbalance between positive and negative classes, the maximum number of training inputs for each class was reduced to 10,000. This utilized all the positive class inputs while randomly subsampling inputs from the negative class. Following this, there were 5,763 “clear” and 1,253 “faint” positive class spectrograms as well as 10,000 negative class spectrograms.

#### 2. Data augmentation

Despite manually-annotated calls showing a high variance in background noise, call duration and note length, for example, data augmentation was applied to further improve input variance. To do this, several pre-defined augmentations supported by Koogu were applied both on waveforms before conversion into spectrograms, and on the spectrograms themselves. These were performed at each epoch, i.e., each time the model passed through the entire training dataset, during the feeding of inputs into the model. This meant that the same original sample could have different levels of augmentations between epochs.

Firstly, Gaussian noise (Schlüter and Grill 2015) was added to 25% of the training input’s waveforms at each epoch to simulate varying levels of background noise. The amount of noise added randomly varied from -20dB to -30dB below the peak dB of the input signal.

The spectrogram was then smeared and squished along the time axis (cf. Madhusudhana 2023). These augmentations were independently added to 20% of the input spectrograms each epoch, at a magnitude of -1,1 (smearing backwards and forwards by one frame of the spectrogram) and -2,2 (stretching and squishing over up to 2 frames of the spectrogram). This process essentially blurred inputs along the time axis, as while the target calls were contained within a standard frequency range, the duration of the signals was highly variable.

Finally, Koogu’s “AlterDistance” augmentation was applied to 25% of the spectrogram inputs. This aimed to mimic the effect of increasing or reducing the distance between the calling gibbon and the receiver, by attenuating or amplifying higher frequencies while keeping lower frequencies relatively unchanged. This was applied by a random factor between -5dB (attenuation) and 5dB (amplification).

#### 3. Network parameters and training

The DenseNet architecture was chosen as the base CNN architecture for this study as it has been shown to achieve comparatively high accuracy with fewer parameters, making it efficient in terms of computational resources (Huang et al. 2016). Early variations of the model suffered from overfitting, occurring when the model learns noise or random fluctuations in the training data rather than the underlying pattern itself. This occurs when the model is too complex relative to its intended task. Bearing this in mind, the standard DenseNet architecture was adapted to a “quasi-DenseNet” architecture (Madhusudhana et al. 2021) which reduces the number of connections within each dense block, limiting model size and complexity. To limit the model’s complexity further and improve computational efficiency, bottleneck layers were also added (cf. Huang et al. 2016). Finally, batch normalization was enabled to improve model convergence. For the final model architecture, see Figure 3.

**FIGURE 3.**
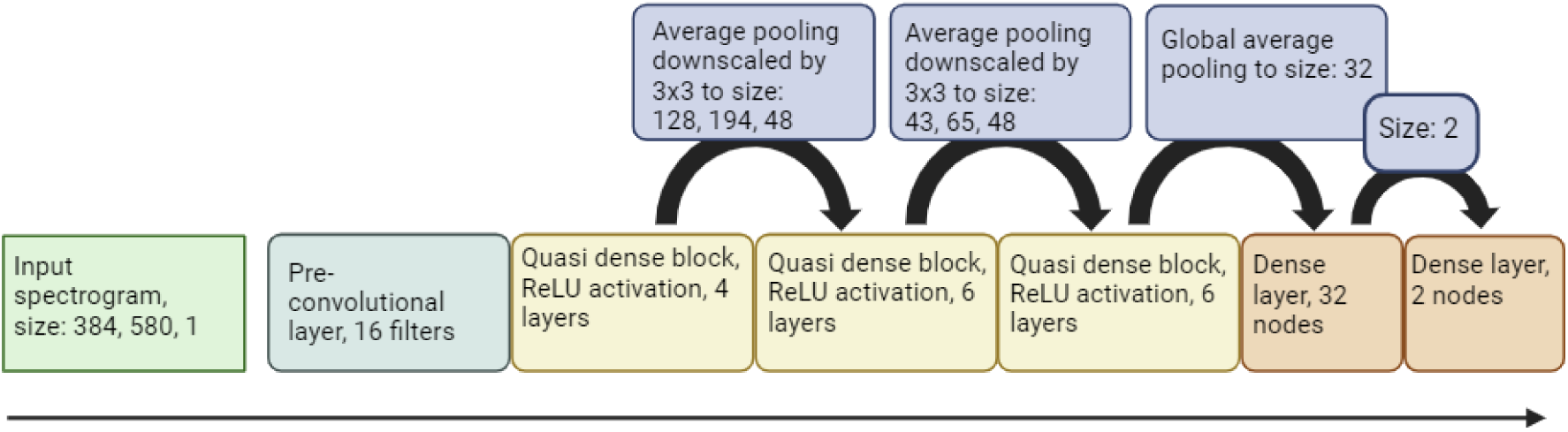
Flowchart showing the final model architecture. The final model had a growth rate of 12, began with a 16-filter pre-convolutional layer, contained 3 quasi-dense blocks with 4, 6, and 6 layers respectively, and finished on a 32-node dense layer. Average pooling layers downscaled the inputs by a factor of 3x3 (height x width) in the transition blocks between quasi-dense blocks. Global Average pooling was used to reduce the spatial dimensions of outputs of the final block to the 32-node feature vectors.

Training inputs were then divided further, with 15% randomly selected as a validation set to evaluate the model’s performance throughout the training process. Dropout layers were added (Srivastava et al. 2014) at a rate of 5% to further reduce overfitting and improve generalization. The models were then trained over 80 epochs using the Adam optimizer (Kingma and Ba 2014) with a minibatch size of 24. The learning rate was initially set at 0.01 and then reduced successively by a factor of 10 at epochs 20, and 40.

#### 4. Testing

Trained models were then applied to the test dataset to provide a preliminary assessment of model performance and establish a desirable detector threshold value. To do this, each test segment was assigned a confidence score by the model between 0 (lowest) and 1 (highest) indicating how likely it was to contain a great call. Performance scores were outputted for thresholds at an interval of 0.01, calculating the number of true positives (TP), false positives (FP) and false negatives (FN). If there was a 100% overlap between a segment and an annotated great call, and the confidence score was above the threshold it was marked as a TP. If there was no- or partial overlap and the score was above the threshold it was marked as an FP. If there was full overlap but the confidence score was below the threshold, then it would be marked as a FN.

These quantities were used to compute recall, Eq. (1), precision, Eq. (2), and F score, Eq. (3), at each threshold (where P = precision and R = recall).

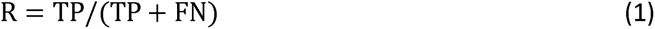

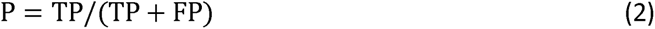

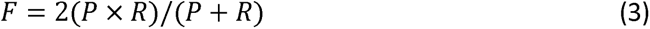

The optimal threshold was then selected using maximum F score, as is a good indicator of overall model performance (Clink et al. 2023).

Once a threshold had been decided, the model was re-run on the test dataset audio files to produce detections and analyse the models’ output. Neighbouring segments for which scores were above the identified threshold were grouped to form a single detection. The score of each detection was then set as the maximum of its component segment scores, and its start and end times were set to the start time of the first segment and the end time of the last segment respectively. These detections were then outputted in the format of Raven Pro selection tables.

### D. Post-processing

Initial inspection of the models’ outputs showed that they produced clusters of detections for each great call, overestimating the number of calls within the data. Detections that overlapped with the first detection in each cluster were grouped together (see Supplementary Material B). These groups were then filtered to retain only the highest-scoring detection(s) within each group to minimize duplicate detections for single great call events. Where the start times of the remaining selections were the same, only one detection was retained. There was no further judication between remaining detections where their scores were tied, as this could indicate that two great calls overlapped or occurred close in time.

Finally, the non-post-processed and post-processed model outputs of the best performing model were evaluated against the manually-annotated test dataset. To do this, we compared the distribution of total detections every 5 minutes across the 4-10 am period using a Kolmogorov-Smirnov test. By comparing the non-post-processed and post-processed model outputs, this also served to validate the effectiveness of the post-processing stage.

## III. RESULTS

### A. Preliminary assessment

The preliminary assessment of the best performing model, trained on only “clear” calls, showed precision ranging from 0.19 to 0.97 and recall ranging from 0.93 to 0.23 at thresholds from 0.01 to 0.99 (Figure 4). This gave a maximum F score of 0.75 at a threshold of 0.36 (precision: 0.80, recall: 0.70).

**FIGURE 4.**
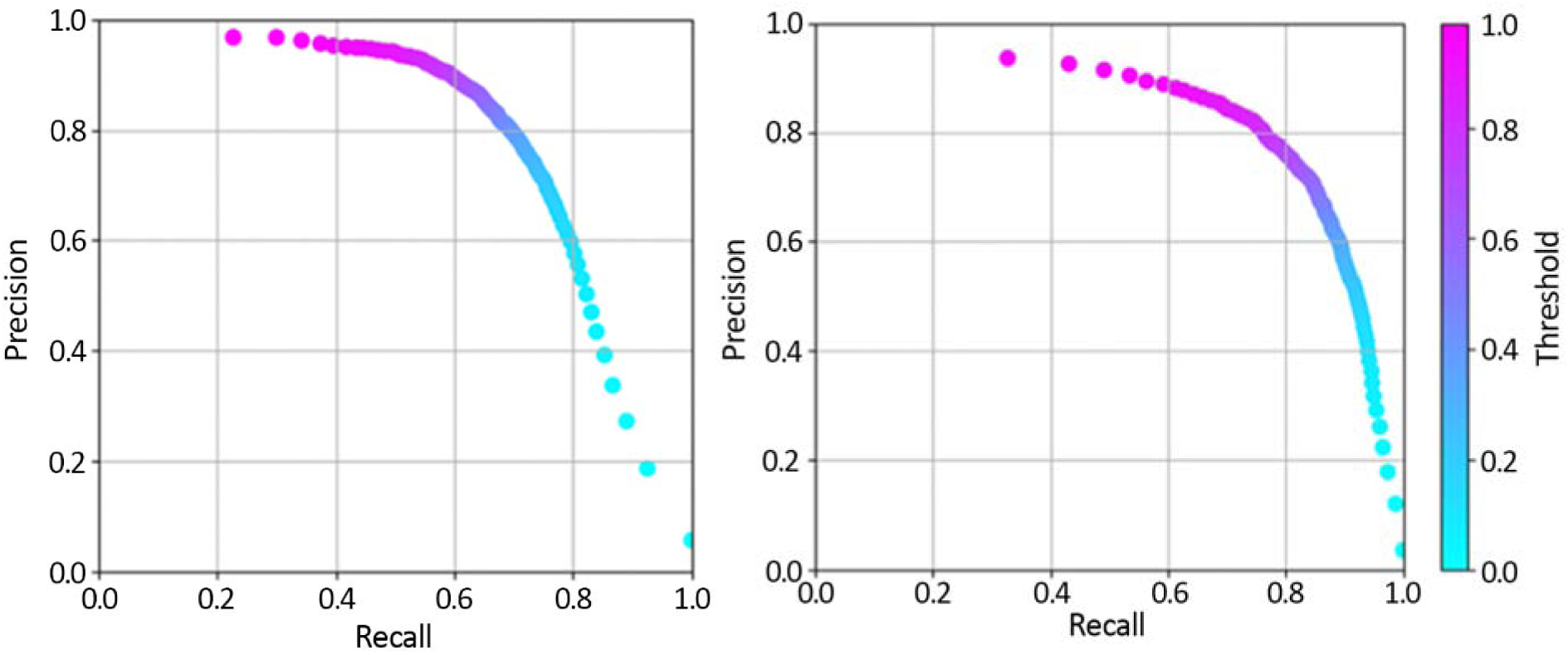
Precision-recall curves of the best performing model tested against all calls (left) and both “clear” and “faint” calls (right).

As the aim was to optimize the model for “clear” and “faint” calls, the testing was re-run excluding “very faint” calls. In this case, precision ranged from 0.12 to 0.94 and recall ranged from 0.99 to 0.33 at thresholds from 0.01 to 0.99 (Figure 4). The maximum F score was improved to 0.78 at a threshold of 0.78 (precision: 0.80, recall: 0.76). To maximize performance for “clear” and “faint” calls while minimizing false positives, a threshold of 0.78 was therefore chosen to evaluate the model’s performance on the test dataset.

### B. Comparison with manual annotations

After re-running the model selected in the preliminary assessment on the test dataset at the desired threshold and processing the output, it produced 535 detections. These were then visually analysed in Raven Pro and compared to the manually-annotated dataset to discern the occurrences of TPs, FPs, and FNs. In this case, a TP instance was defined as any model detection that overlapped with a great call. The model was found to have produced 511 TPs and 24 FPs, missing a further 133 calls (FNs). This gives a precision of 0.96, a recall of 0.79, and an F score of 0.87.

Upon closer inspection of the post-processed model output, 86 of the TP detections were the result of repeated detections for singular great call events. Additionally, 22 FNs were classed as TPs before the post-processing stage. These were created as a result of inadvertently grouping detections where two great calls overlapped in time or were adjacent to one another.

Overall, the non-post-processed model identified 409 (78%) of the 522 manually-annotated calls. This included 98% of all “clear” calls, 73% of “faint” calls, and 44% of “very faint” calls. Out of the FN instances before post-processing, 77% were “very faint” calls with only 0.05% representing 6 missed “clear” annotations. A further 38 great calls were identified which had been missed in the manual annotation stage, including 6 “clear”, 8 “faint” and 24 “very faint” calls.

The distribution of total detections every 5 minutes across the 4-10 am period for both the non-post-processed and post-processed model outputs were compared against the manually-annotated test dataset (Figure 5). Kolmogorov–Smirnov tests indicated no significant difference (D = 0.036, p > 0.05 and D = 0.045, p > 0.05 respectively). Also, we found no significant difference between pre-processed and post-processed data (D = 0.020, p > 0.05).

**FIGURE 5.**
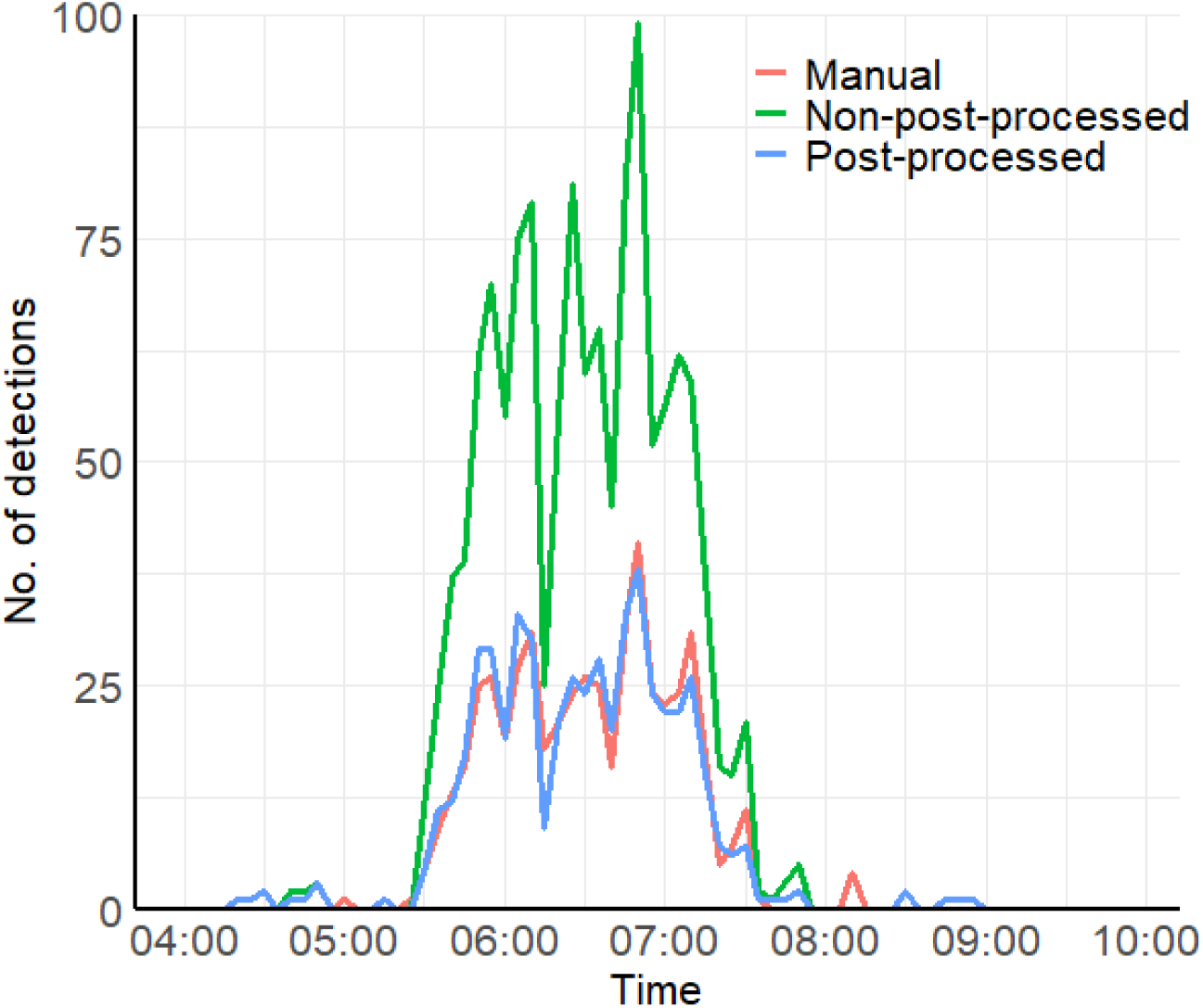
Histogram showing the number of great calls detected every 5 minutes between 04:00 and 10:00 for the manually annotated test dataset, as well as the output for the non-post-processed model, and the post-processed model when ran on the test dataset.

## IV. DISCUSSION

Our best performing model was effective at detecting high-quality *H. albibarbis* great calls with a low rate of false positives. The best performing model (F score: 0.87) exceeded previously reported SVM models for detecting gibbon vocalisations e.g., *Hylobates funereus* detector, F score: 0.78 (Clink et al. 2023) and was largely comparable to other CNN models for gibbon great calls, e.g., *Nomascus hainanus* detector, F score: 0.91 (Dufourq et al. 2021), *Nomascus concolor* DenseNet detector, F score: 0.92 (Zhou et al. 2023).

We found that the best performing model detected 38 great calls that had been missed during the manual annotation stage, amounting to 6.8% of the total number of target events. Human listening is subject to error (Brauer et al. 2016; Knight et al. 2017), and this study relied on multiple human observers with differing levels of training to construct the manually-annotated dataset. While it has been shown that human observers with less experience may perform worse than some automated detectors (Jennings et al. 2008), the consensus from multiple observers may have reduced the level of human error (Drake et al. 2016). Furthermore, signals with low signal to noise ratio (SNR) are difficult for both humans and machine-learning algorithms to detect (Knight et al. 2017). Precision and recall are often measured relative to manually-annotated datasets, but these are not always perfect. With this in mind, it has been recommended to view the manually-annotated dataset as the output from an alternative detector rather than a ground-truth set (Knight et al. 2017).

Due to the apparent likelihood of great calls to be missed by manual annotation (Knight et al. 2017), it is unrealistic to presume that all target calls were identified in the 210-hour training dataset. This could pose a problem when training the model if any of the randomly selected unannotated time periods for the negative class contained target events, potentially increasing the rate of false negatives. Dufourq et al. (2021) found that better results were obtained by specifically including negative class segments with typical ambient noise, such as other species’ vocalisations, which could potentially confuse the classifier. This method of ‘hand picking’ the negative class could reduce the false negative rate and the false positive rate by negatively labelling potentially confusing information. The low false-positive rate reported in this study, as well as the low false-negative rate for “clear” calls, suggests that our method of randomly selecting the negative class was sufficient for our aims. Where reducing the false-negative rate has greater importance, such as when identifying infrequent vocalisations, a more thorough approach could be preferable. Nonetheless, it is important to recognize the trade-off between ensuring that the negative class contains as little erroneous information as possible and the time required to construct an adequate training dataset.

In this study, we categorized selected calls according to their quality (cf. Cañas et al. 2023), although we acknowledge that the distinction of “clear”, “faint” and “very faint” great-call selections was not based on p measurements. These categories cannot be translated directly to distance; however, in most cases, it is likely that “clear” calls were recorded from gibbons singing closer to the ARUs. Overall, our best performing model identified 98% of all “clear” calls, and only missed 6 from the manually-annotated dataset. This was comparable to the human observers, who also missed 6 “clear” calls. The model performed worse than the human detector at detecting “faint” and “very faint” calls, however, picking up 73% and 44% of the manually-annotated instances, respectively. With this in mind, the results suggest that detection likelihood is affected by the caller-ARU distance (Spillmann et al. 2015). This supports the suggestion by Jahn et al. (2017) that the difference in recall between an automated detector and a human listener is caused by the former having a smaller detection radius. Future studies should aim to apply relationships between signal strength and the distance of the source from the receiver to estimate call detection probability over distance. This will help to determine the effective area being sampled by PAM studies.

Although the model did not have perfect recall, the importance of detecting all great calls within a recording will depend on the research question. With regards to calling activity over time, the distribution of calling frequency detected by the model was not significantly different to that of the manual annotations. Therefore, our model can be used to reliably estimate spatiotemporal variations in *H. albibarbis* calling activity. Through analysing recordings from multiple habitats, our model could be applied to understand the relative importance of forest subtypes for the species, as singing behaviour is density-dependent (Cheyne et al. 2008), with less singing activity at lower group densities. This could operate on a continuous long-term time frame which would be hard to achieve when relying on human observers in the field.

For an in-depth understanding of gibbon group abundance and density, further information is necessary. One method is to use individuals as the sampling unit (Buckland 2006) by analysing their call structure (Clink et al. 2023), or by localizing vocalisations using estimates of direction to the source from multiple ARUs (Stevenson et al. 2015). This may prove highly effective at estimating gibbon abundance and density over the short term yet could prove too complex over the course of hundreds, or thousands of hours of audio. An alternative method is to estimate vocalisation density per unit time, apply an estimation of vocalisation rate, and then convert vocalisation density into group density (Marques et al. 2013). This does require a knowledge of the area covered by each ARU, yet, for this task, Marques et al. (2013) notes that automated detectors need not perform extraordinarily well so long as true positive and false positive rates are characterized accurately. This method does not require for the effective area of ARUs to overlap, so in theory, could monitor a larger area with the same resources.

While post-processing greatly reduced the number of repeated detections for single great call events, these would still account for many false positives if only one true positive per great call is allowed. The post-processing protocol did not seek to adjudicate between detections if there was a tie in the highest scoring instances within a group, as in some cases this represented two great call events close in time. In fact, 22 false negatives derived from instances when the grouping of detections failed to take this into account, and limiting the number of detections per group to one would have increased the false negative rate further. An alternative approach would be an improved post-processing stage to better interpret the output of the model. In audio classification, CNNs inspect audio recordings as image-like segments and so are unable to use broader-scale contextual information, such as whether the current point in a recording is preceded by a target call (Wang et al. 2022). It has therefore been proposed to combine CNNs with other machine learning techniques, such as Hidden Markov Models (HMMs) or deep learning techniques, such as Recurrent Neural Networks (RNNs). Postprocessing the output of a CNN with a HMM or an RNN has been shown to improve the F score (Madhusudhana et al. 2021; Wang et al. 2022). For instance, Wang et al. (2022) showed that the application of a combined Convolutional Recurrent Neural Network (CRNN) reduced error rates arising from overestimation of gibbon calls (49-54%) to 0.5%. It was noted, however, that while post-processing the model output with a HMM performed second best, it required much less computational power. Future work should therefore consider both of these options when attempting to improve on our post-processing method.

A key aim of this study was to help address the analysis bottleneck evident in PAM data by improving on the time taken to manually analyse recordings. During the batch processing stage, it took 19 seconds for our trained model to process each hour of test recordings. This greatly improved on a human processing speed of minimum 1 hour per hour of audio for this study, varying depending on the level of observer experience. One caveat is the significant amount of time required to construct a manually-annotated dataset to train and test a CNN when compared to other machine learning approaches (Stowell 2022). Despite our study showing how data augmentation can be effective where training data is limited, careful consideration should be taken when under time-pressure if no training datasets are already available. In some cases, it may be better to adopt an approach that requires less data to develop, such as an SVM or a GMM (Clink et al. 2023).

Finally, Stowell (2022) noted that it was increasingly common to evaluate deep learning models on test sets specifically designed to differ in some respects from the training data, such as location, signal to noise ratio, or by season. In this case, the model was designed with application to the MBERF bioacoustic dataset in mind, and so it is appropriate that the test data came from the same location as the training data. Test inputs spanned across all times of the year and were recorded from three different habitats. This ensured a variety of potential sound environments were included in the testing stage. However, for applications in other locations, especially outside the Rungan Forest Landscape, it would be advantageous to first test the model on recordings captured elsewhere within *H. albibarbis’* range.

## V. CONCLUSION

Our study demonstrates how an open-source deep-learning framework can be adapted to produce a CNN capable of detecting *H. albibarbis* great calls, performing at a comparable level to similar CNN approaches for gibbon great calls. Our model performed best on the highest-quality calls and yielded a low false-positive rate, meeting the objectives of this study. There was a much lower likelihood of successful detection for the lowest-quality calls, however, and future studies should aim to estimate call detection probability over distance to determine the effective area being sampled.

Further development of the post-processing stage could help to reduce the number of repeat detections for each call. However, the current output can be used to estimate calling rate over time. Further work should seek to apply this model to long-term acoustic datasets over a variety of habitats to study spatial and temporal variation in gibbon calling activity. Furthermore, in combination with future studies on sound propagation of gibbon vocalisations, this represents an opportunity to monitor *H. albibarbis’* populations on an ever-greater spatiotemporal scale. Our work presents some key considerations to inform decision-making for such projects, and a full workflow script to visualize how these can be implemented in developing an automated detector.

## SUPPLEMENTARY MATERIAL

For the full Koogu workflow, see Supplementary Material A (SuppPub1.txt). For the post-processing script, see Supplementary Material B (SuppPub2.txt).

## Supporting information

SuppPub1

SuppPub2

## ACKNOWLEDGEMENTS

We thank the Universitas Muhammadiyah Palangkaraya and Borneo Nature Foundation for access to and support in the MBERF. We thank the Indonesian government for permission to carry out this research (RISTEK-DIKTI permit #189/SIP/FRP/E5/Dit.KI/VI/2018 and 43/E5/E5.4/SIP.EXT/2019 [Wendy M. Erb], BRIN foreign research permits 223/SIP/IV/FR/5/2023 [F. J. F. van Veen] and 225/SIP/IV/FR/5/2023 [A. F. Owens]). We thank Erik Estrada and Rido for their invaluable support in deploying and maintaining the ARUs and data in the field. We also thank Georgia Allen, Amy Barron, Sophie Carpenter and Elena Gough for their contributions in manually annotating the dataset. In addition, we acknowledge the contributions of the Indigenous Dayak Ngaju community of Mungku Baru, including Pak Edo, Pak Yuli, Pak Viktor and Rico, who supported the data collection and shared their knowledge together with local conservation practitioners from Yayasan Borneo Nature. This project has been a collaboration between international and Indonesian researchers from the start and this is recognized in joint authorship here. Finally, please note the sponsors of this research, awarded to WME: Primate Conservation, Incorporated; British Academy; Conservation International; American Association of Physical Anthropologists; International Primatological Society; and American Institute for Indonesian Studies.

AFO, FJFvV, WME, MAI and KH developed the project and concept; M, TMS and S Maimunah provided permissions and guidance for field study design and execution; AFO, S Madusudhana and MS carried out building and analysis of the model; AFO drafted the manuscript to which all other authors then contributed to produce the final version.

## AUTHOUR DECLARATIONS

The authors declare no competing interests.

## DATA AVAILABILITY

Recordings of great calls for both the training and testing datasets are openly available on Zenodo at 10.5281/zenodo.10926304.

## ETHICS APPROVAL

Ethical approval was provided by the University of Exeter (application ID: 1845574), BRIN (application number: 22022023000026) and Institutional Animal Care and Use Committee of Rutgers, the State University of New Jersey protocol number: PROTO201800073.

