## Supplementary material for "Automated detection of Bornean white-bearded gibbon (*Hylobates albibarbis*) vocalisations using an open-source framework for deep learning": SuppPub2

#Code to process output from model, removing repeat detections

# Initialize variables
i = 1
max_grp = 1
recent_grp = 1

#Start/end times of first detection
first_grp_st = df$st[1]
first_grp_fin = df$fin[1]

# Loop starts here, iterate over each row
while (i <= nrow(df)) {
  if (i == 1) {
    df$group[i] = 1
  } else {
    # Compare the start time of the current row to the start and end times of the first row within the most recent group
    # Is it between the start and end times of the row above
    if (df$st[i] >= first_grp_st && df$st[i] <= first_grp_fin) {
      df$group[i] = recent_grp
    } else {
      recent_grp = max_grp + 1 # Establish new group number
      df$group[i] = recent_grp #Assign new group number to row
      max_grp = recent_grp #Track highest group number assignment as of yet
      
      # Update the start and finish times for the new group
      first_grp_st = df$st[i]
      first_grp_fin = df$fin[i]
    }
  }
  
  i = i + 1
}

#Filter by highest scoring detection in each group
post_df <- df %>%
  group_by(group) %>%
  filter(Score == max(Score)) %>%
  ungroup()

#Remove duplicates when start time is the same
post_df <- post_df %>%
  group_by(group, st) %>%
  filter(row_number() == 1) %>%
  ungroup()
